# A reduced order model of the spine to study pediatric scoliosis

**DOI:** 10.1101/2020.04.20.051995

**Authors:** Sunder Neelakantan, Prashant K. Purohit, Saba Pasha

## Abstract

The S-shaped curvature of the spine has been hypothesized as the underlying mechanical cause of adolescent idiopathic scoliosis. In earlier work we proposed a reduced order model in which the spine was viewed as an S-shaped elastic rod under torsion and bending. Here, we simulate the deformation of S-shaped rods of a wide range of curvatures and inflection points under a fixed mechanical loading. Our analysis determines three distinct axial projection patterns of these S-shaped rods: two loop (in opposite directions) patterns and one lemniscate pattern. We further identify the curve characteristics associated with each deformation pattern showing that for rods deforming in a loop 1 shape the position of the inflection point is the highest and the curvature of the rod is smaller compared to the other two types. For rods deforming in the loop 2 shape the position of the inflection point is the lowest (closer to the fixed base) and the curvatures are higher than the other two types. These patterns matched the common clinically observed scoliotic curves - Lenke 1 and Lenke 5. Our elastic rod model predicts deformations that are similar to those of a pediatric spine and it can differentiate between the clinically observed deformation patterns. This provides validation to the hypothesis that changes in the sagittal profile of the spine can be a mechanical factor in parthenogenesis of pediatric idiopathic scoliosis.

## 1 Introduction

During the fast growth period around puberty, some pediatric spines deform in three dimensions leading to scoliosis[5, 6, 7, 30]. While the pathogenesis of this disease remains unknown[12, 16], the side view S-shaped curvature of the fast growing, flexible, immature, slender spines has been hypothesized as an underlying mechanical cause of adolescent scoliosis[8, 21, 26, 28]. It has been clinically shown that at an early stage of scoliosis, the sagittal curvature of the spine is different between scoliotic and non-scoliotic subjects of similar age and sex [29]. However, as the scoliosis changes the spinal alignment in three-dimensions[4], even at an early stage of the disease,it is challenging to evaluate the role of the true sagittal alignment of the spine in induction of scoliosis. As the idiopathic scoliotic patients are otherwise healthy, identifying these patients before the onset of scoliosis in order to obtain their patterns of sagittal profile is difficult[7]. Hence we turn to simulations to test the hypothesis that there is an association between the sagittal curvature and deformity patterns of the spine[3, 14, 17].

A reduced order model of the pediatric spine for studying scoliosis was proposed by Pasha[21]. Using finite element analysis, this model simulates the spine as an S-shaped slender elastic rod and applies gravity to simulate the body weight and a torsion along the spine to simulate the body mass asymmetry [3, 10, 13, 23]. In the current study, we wish to determine the geometrical parameters that affect the deformation of such slender elastic rods while mechanical loading remains unchanged. One way to do this is by finite element simulation as was done previously by modeling the spine as a linear elastic material[21]. Repeating such finite element simulations while iterating over various initial shapes obtained by permutation of geometric parameters is computationally prohibitive. To reduce computational cost, we developed a semi-analytic model using Kirchhoff equations[18] to solve for the deformation given the initial geometry and loading on an S-shaped elastic rod[19]. The analytical model allows us to compute the deformed configuration of a rod subject to loads through a set of ordinary differential equations. The reduced computational complexity allows us to iterate over all permutations of the geometric parameters describing the sagittal shape of the spine.

To this end, we simulate the curvature of the spine by two regions of constant curvatures. We define the sagittal profile of the spine using 5 parameters shown in fig.(1) which allows us to vary the shape of the sagittal curve systematically. Using this simplified geometry, we hypothesize that under the same mechanical loading and mechanical properties, the geometric parameters – the two curvatures of the S-shaped rod, position of the inflection point, and the slope of the curve at the lowest point with respect to the horizontal axis – can significantly impact the deformation of the rod.

**Fig. 1:**
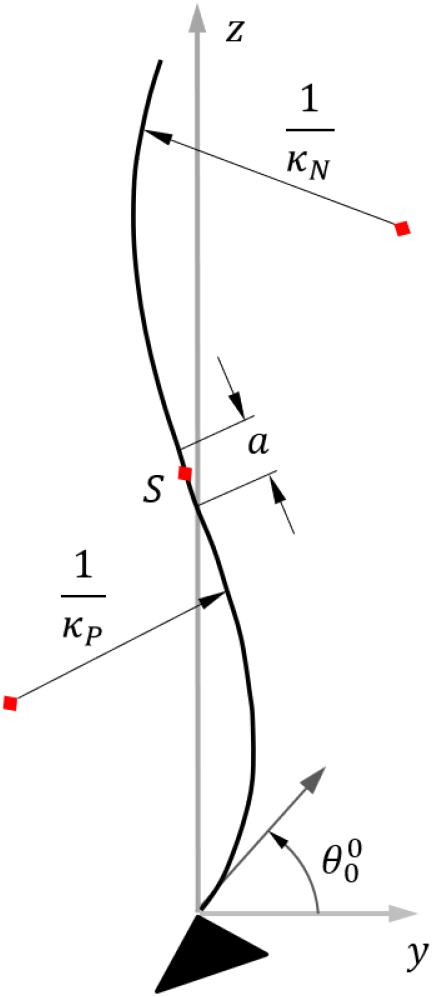
Schematic of the rod. *α*_0_ + *θ*_0_ = *π*/2 and *ϕ*_0_ = *π*/2

## 2 Methods

Here we describe the elastic rod model for the spine to investigate the effects of the different geometric parameters. Since our goal is to focus on the geometric parameters of the sagittal curvature, we we will hold the bending modulus of our S-shaped rod fixed for all the simulations. We present a schematic in fig.(1) to label the geometric parameters

The spine is modeled as an S-shaped elastic rod. The undeformed spine is assumed to rest on the **y** − **z** plane with the base of the spine at the origin in the lab coordinate system given by [**e**_*x*_ **e**_*y*_ **e**_*z*_]. We define an arc length coordinate along the center-line s; a point located at *s* in the reference configuration moves in the deformed configuration to **r**(*s*) = *x*(*s*)**e**_*x*_ + *y*(*s*)**e**_*y*_ + *z*(*s*)**e**_*z*_. The rod is then subject to a body force along the −*υe z* direction which is caused due to the weight distribution of the upper body. It is also subject to moments at the ends to simulate asymmetry in the body mass[23].

The Frenet-Serret frame for the rod is 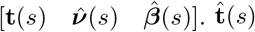 is the tangent vector, 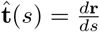. The rod is assumed in-extensible, hence 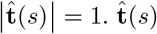 can be expressed in the lab coordinate system as:

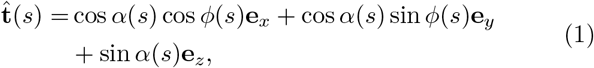

where *α* is the polar angle measured from the *x* − *y* plane and *ϕ* is the azimuthal angle used in conventional spherical polar coordinates. 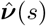 and 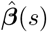 are the normal and binormal vectors, respectively, and they are computed using the Frenet-Serret equations:

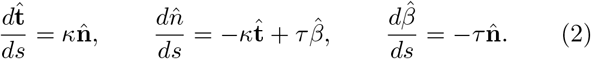

The curvature *κ*(*s*) and the torsion *τ*(*s*) of the rod are obtained from the above equations[20]. This completes the kinematic description of the rod.

### 2.0.1 Mechanics

Next, we present the equilibrium equations for the rod. We begin with the conservation of linear momentum[1, 2],

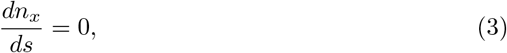

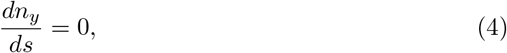

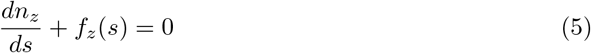

where **n**(*s*) = [*n_x_*(*s*) *n_y_*(*s*) *n_z_*(*s*)] is the internal force in the rod. **f** = *f_z_*(*s*)**e**_*z*_ is the body force on the rod and is directed only along the **e**_*z*_ direction due to gravity. For *f_z_*(*s*), we use the values found in Pasha et al.[23]. The rod is assumed in-extensible, hence there is no constitutive law for **n**(*s*). **n**(*s*) must be determined as part of the solution of the boundary value problem for the rod. We use eq.(2) and (3) to get

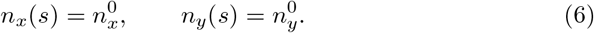

Since there are no forces in the **e**_*x*_ and **e**_*y*_ directions, 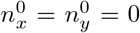. The conservation of angular momentum of the rod states that

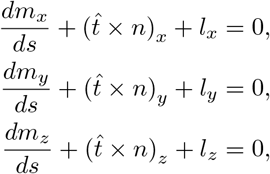

where **m**(*s*) = [*m_x_*(*s*) *m_y_*(*s*) *m_z_*(*s*)] is the internal moment in the rod in the lab frame and **l** is the body moment per unit length. We set **1** = **0** in this work. Since *n_x_* = *n_y_* = 0, the balance of angular momentum becomes:

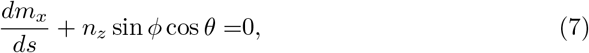

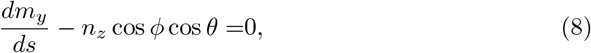

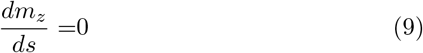

so we get *m_z_* = *T* from eq.(9), a constant which we compute from torque boundary condition applied at *s* = 0.

The internal moment can be written in the Frenet frame as 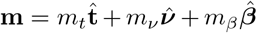 which is convenient if we use the following simple constitutive relation:

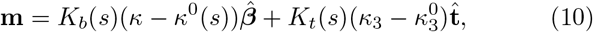

where *K_b_*(*s*) and *K_t_*(*s*) are the bending and twisting moduli of the elastic rod, respectively. The curvature functions in the stress free state are given by *κ*^0^(*s*) and 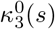, respectively. We assume 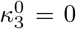. This constitutive law is a special case of a general form given by 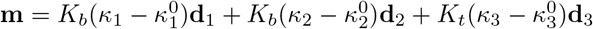 where [**d**_1_(*s*) **d**_2_(*s*) **d**_3_(*s*)] is a material frame that convects with the arc-length coordinate *s*.

Plugging this constitutive law into the conservation of angular momentum and considering the component along the 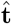 direction only gives[1, 2]

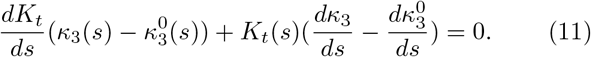

If we define 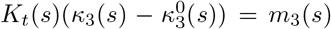, then eqn. (11) shows that 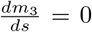, or *m*_3_ = *const*. We also set *K_b_*(*s*) as constant since our focus is on the effects of the geometric parameters. Then, from the constitutive law, we can write the moment in the lab frame as

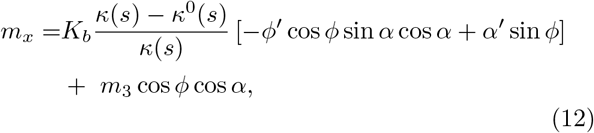

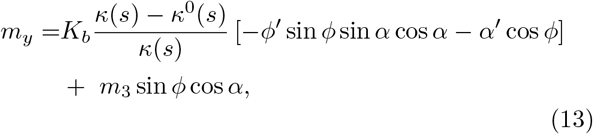

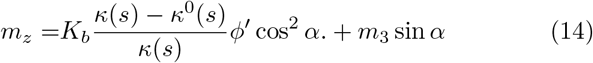

The elimination *K_b_, κ*^0^(*s*) and *κ*(*s*) gives:

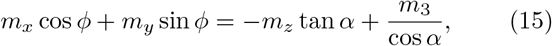

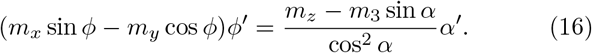

Then, we solve Eqn. (15) to get

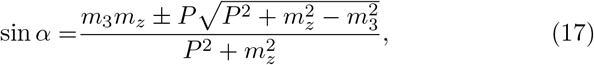

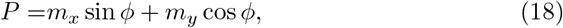

where we determine the solution branch from *α*_0_. We compute *ϕ*′(*s*) using eqn.(16) to get

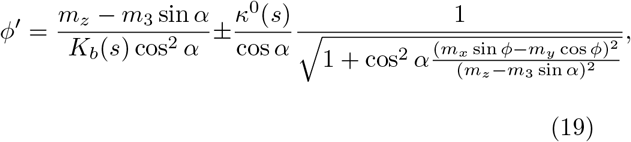

where the ± sign is dependent on the sign of *ϕ*′(*s*).

Finally, the deformed curve can be determined using

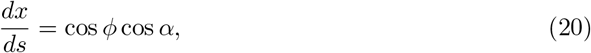

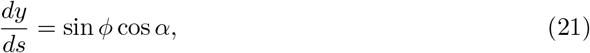

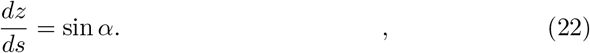

This along with eq.(5), (7), (8) and (19) form the governing equations of the system along with the boundary conditions given by:

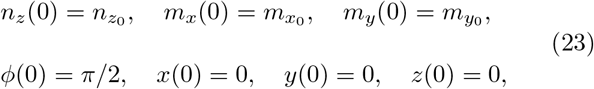

where *n*_*z*0_ is the weight of the upper body and *m*_*x*_0__ and *m*_*y*_0__ are used to account for the loading asymmetry experienced by a scoliotic spine.

### 2.1 Phase plot

Now, we will use the model above to study the effects of the different geometrical parameters of the curved rod. *K_b_* is set to an arbitrary constant. We will simplify *κ*^0^(*s*) using the following function:

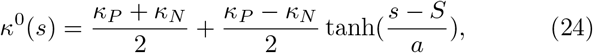

where *S* is the point of inflection of the rod and *a* is the distance over which the curvature of the rod goes from *κ_P_* to *κ_N_ · κ_N_/κ_P_* are the curvatures of the lower/upper portion of the rod (above and below the inflection point respectively).

We will focus on the effects of varying *κ_P_, κ_N_, S, a* and *θ*_0_. The range of values for these parameters used in this study are in table.(1). To reduce the data space, we study the parameters that cause the largest change in deformations. We restrict *θ*_0_ to the three major values found in clinical studies[24]. The range of values for *κ_P_, κ_N_, S* and *a* are based on maximum and minimum values of these parameters in Pasha et al.[25] who investigated the geometric shapes of spines of patients with adolescent idiopathic scoliosis. For a range of *a* (0.01 - 0.1), we did not observe significant changes in deformation when we varied *a* over this range. So, we keep *a* fixed in all our computations and focus on the remaining four geometric parameters. As for the loading and mechanical properties, since the deformation in the axial view is dominated by *m_z_* (see the Discussion section), we keep the ratio *m_z_/K_B_* in the range observed in our previous study[19].

**Table 1:** Parameter values

| $\kappa_P$ [m<sup>-1</sup>] | $\kappa_N$ [m<sup>-1</sup>] | $S$ [m] | $\theta_0$ [°] | $a$ [m] | $K_b$ [Pa] | $m_z$ [Nm] | $m_3$ [Nm] | $m_x(0)$ [Nm] |
| --- | --- | --- | --- | --- | --- | --- | --- | --- |
| 0.1 - 5 | 0.1 - 5 | 0.3 - 0.8 | 31°, 37°, 44° | 0.01 | 10 | -2 | 0 | -8 |

### 2.2 Uprightness

To ensure the deformed state of the rod is physiologically acceptable, we restrict the values of the parameters such that the initial configuration of the spine remains upright.The shape of the spine is assumed to lie in the **y** − **z** plane in the absence of loads. Hence, when *ϕ*(*s*) = *π*/2, we can compute the initial curve from *κ*^0^(*s*) using the following system of ODEs

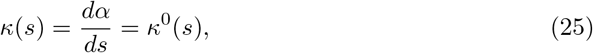

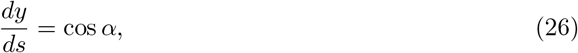

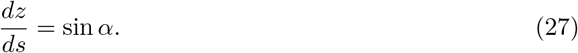

We limit the **y** displacement of the top of the spine to 20% of the **z** displacement in the initial configuration to filter out unrealistic shapes where there is a large displacement between the head-pelvis i.e. the two ends of the rod. This constraint boils down to

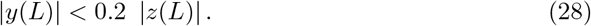

Now we apply the analytical model to find the deformation over the range of parameters. As shown previously by Pasha et al.[25], the most common curve types in scoliosis deform in a loop or lemniscate shape in the axial plane (**x** − **y** plane). We classify the deformed shapes as loop or lemniscate based on the number of points of intersection the curve in the **x** − **y** projection has with the line-segment joining the two end points of the curve. The curve is classified as a loop shape if the line-segment does not intersect the curve and as a lemniscate shape if the line-segment intersects with the curve at least once (we exclude the intersection at the end-points). We perform this check over the range of parameters given in table.(1).

We create a scatter plot of points in the data space which satisfy the lemniscate condition using *κ_P_, κ_N_* and *S* as the axes of the plot. The boundary points are filtered and surfaces are fit to the points. A polynomial fit is used to separate the regions into loop and lemniscate shapes. A similar procedure is followed for points satisfying the upright condition and a polynomial fit is used to create surfaces to separate regions where the undeformed rod is upright from those where it is not.

For each *θ*_0_, we plot all the initial configurations that deform into loop and lemniscate shapes, respectively, along with the average curve for each case. The equations of the surfaces for the lemniscate condition along with the coefficient of determination of the fit (**R^2^**) are given in table.(S1) in the supplementary material. The resulting three surface plots at the three *θ*_0_ values appear in the results section.

## 3 Results

Fig.(2) presents the loop-lemniscate classification regions and the condition of uprightness in the data space. We see in fig.(2a) that two surfaces divide the volume of interest into 3 parts comprising of two loop regions and one lemniscate region. We label the loop region with low *κ_P_* and *S* as “loop2” and the other region as “loop1”. Only variable ranges bounded by both uprightness surfaces and loop-lemniscate classification surfaces are presented in the following sections. The different regions are classified accordingly and serve as a reference when we consider both conditions together in the following plots.

**Fig. 2:**
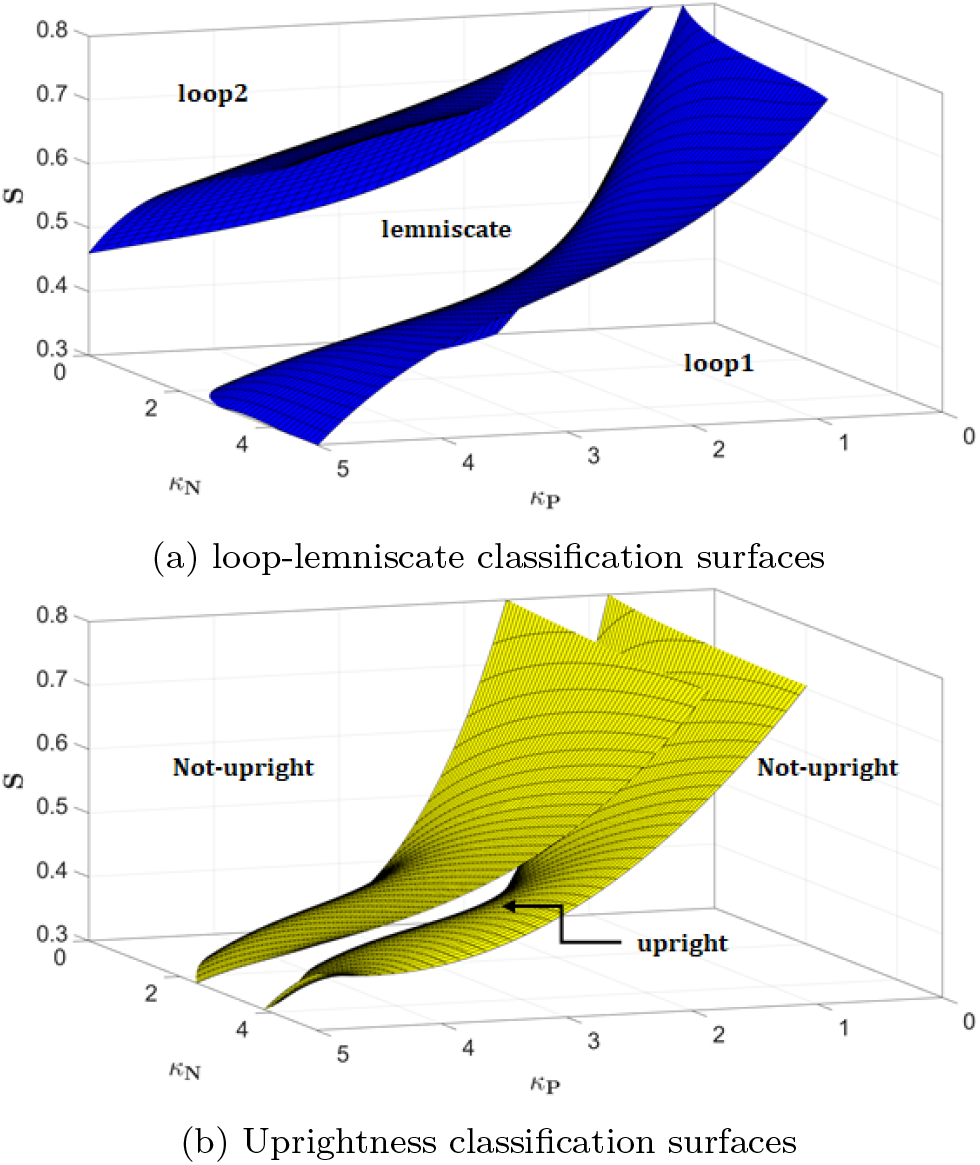
Representative classification surfaces with regions marked

Next, we present the combined surface plots in fig.(3). We present the surface plots for 3 angles - 31°, 37°, 44°. The surfaces span the entire parameter space, so that we can visualize any trends that develop in the region of interest. We have included the equations of the surfaces and the fit information in table.(S1) of the supplementary material. We have also presented the range of values in table.(2). We also present the equations for the uprightness check surface (yellow surface) in table.(S2) of the supplementary material.

**Table 2:** Average and range of the geometrical parameter values describing the sagittal profile for Loop1, Lemniscate, and Loop 2 cases. The parameter values of the average curve is in bold while the range of values is given with [].

| Case | $\theta_0$ | $\kappa_P$ | $\kappa_N$ | $S$ |
| --- | --- | --- | --- | --- |
| Loop1 | 31° | **1.36** [0.85 : 1.60] | **0.32** [0.10 : 0.85] | **0.81** [0.70 : 0.80] |
|  | 37° | **1.50** [0.95 : 1.90] | **0.44** [0.10 : 1.45] | **0.80** [0.65 : 0.80] |
|  | 44° | **1.75** [1.20 : 2.30] | **0.62** [0.10 : 2.05] | **0.80** [0.60 : 0.80] |
| Lemniscate | 31° | **2.06** [0.75 : 5.00] | **1.75** [0.10 : 5.00] | **0.56** [0.30 : 0.80] |
|  | 37° | **2.71** [1.10 : 5.00] | **2.50** [0.45 : 5.00] | **0.55** [0.30 : 0.80] |
|  | 44° | **2.74** [1.25 : 5.00] | **2.12** [0.10 : 5.00] | **0.55** [0.30 : 0.80] |
| Loop2 | 31° | **2.78** [0.85 : 5.00] | **2.72** [0.10 : 5.00] | **0.41** [0.30 : 0.80] |
|  | 37° | **4.35** [3.10 : 5.00] | **4.19** [2.75 : 5.00] | **0.40** [0.30 : 0.45] |
|  | 44° | **4.03** [2.55 : 5.00] | **3.18** [1.05 : 5.00] | **0.40** [0.30 : 0.50] |

**Fig. 3:**
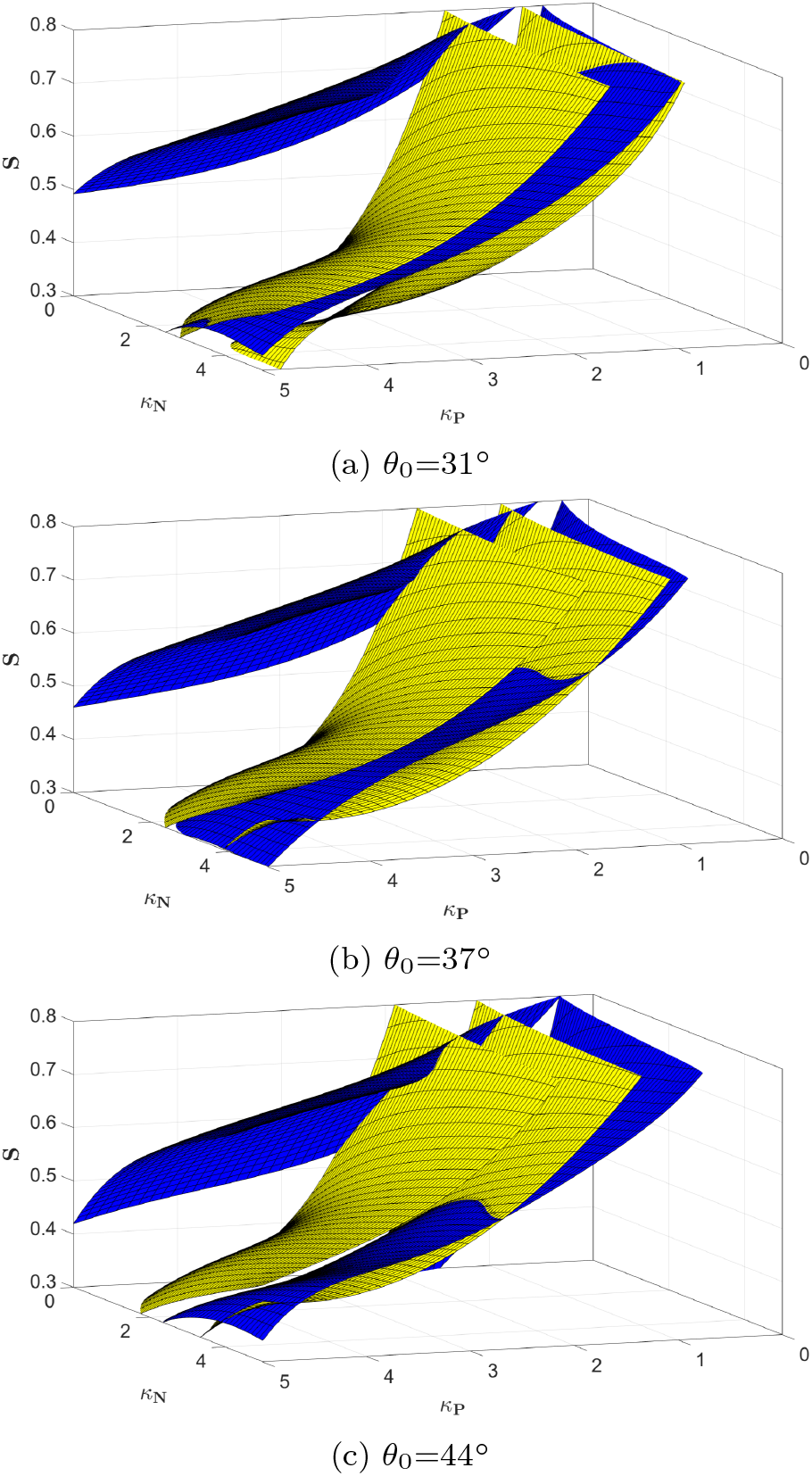
The loop and lemniscate regions for each case of *θ*_0_.

From fig.(2b), we see that the range of admissible *κ_P_* values that satisfy the uprightness condition is smaller than the range of *κ_N_* values for each *S*. Recall that the range of *a* is also very small and it has minimal effect on the classification of deformation patterns.

We present the shape of the initial configuration that lead to loop and lemniscate shapes respectively, in the deformed configuration in fig.(4). The initial shape of curve (black line) along with the range of the curves (shaded area) for each *θ*_0_ are shown in fig.(4). The corresponding sagittal profile values are presented in table.(2). As seen in both fig.(4) and table.(2), the position of the inflection point is the highest, closer to the top, in loop 1 and decreases in lemniscate and loop 2 cases. However, the changes in the inflection point within each case as the *θ*_0_ changes is small. Both *κ_P_* and *κ_N_* are the smallest in loop 1 cases and highest in loop 2 cases, suggesting a more flat sagittal profile in loop 1 patients compared to the lemniscate and loop 2 cases.

**Fig. 4:**
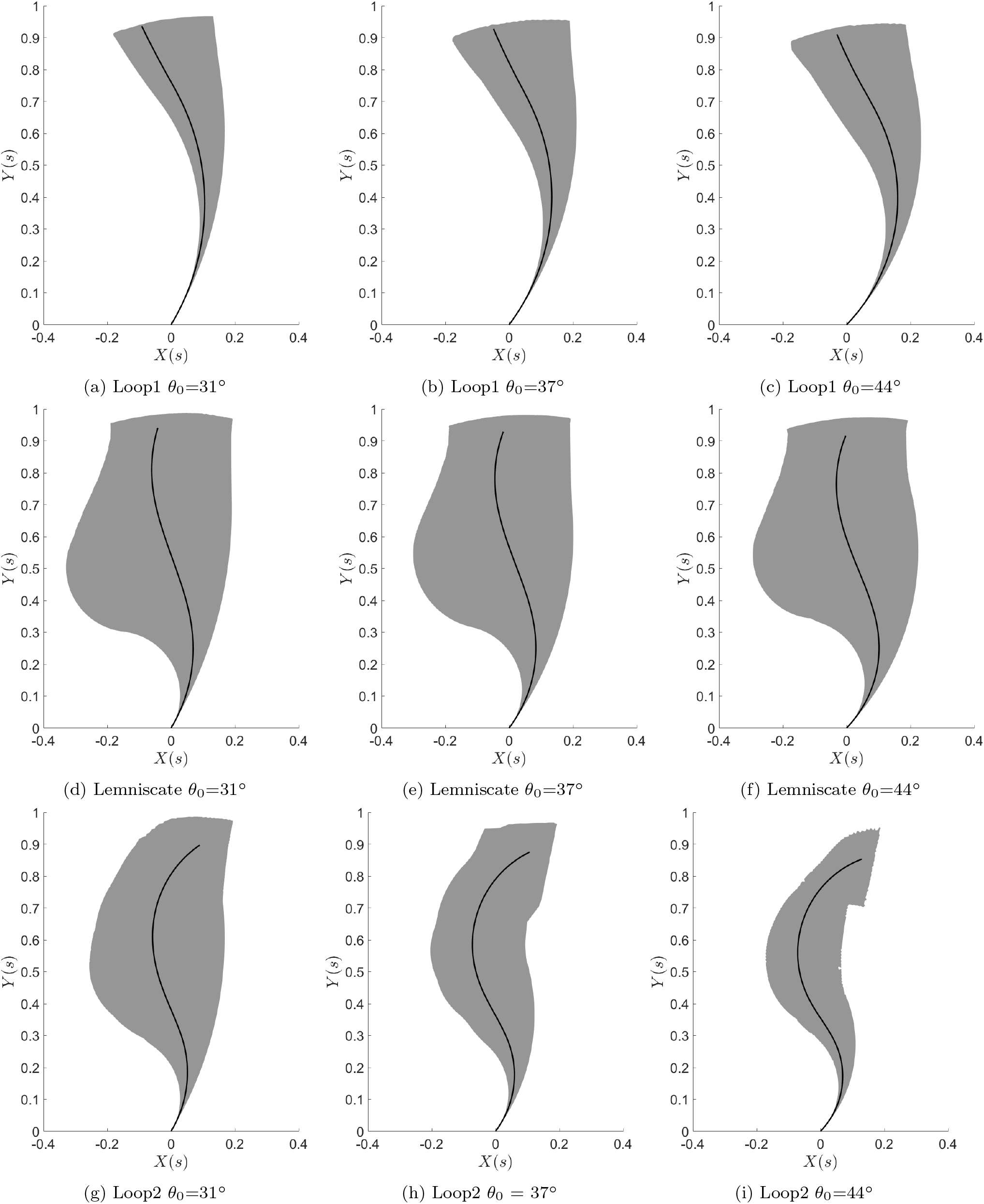
The initial configuration to get loop and lemniscate shapes for each case of *θ*_0_. The line is the average curve and the grey region corresponds to the range.

## 4 Discussion

We analyzed the deformation patterns of S-shaped elastic rods as it pertains to the pediatric spine to investigate the role of the geometrical parameters of the sagittal spine in induction of scoliosis. Our results determined, under physiological conditions, an S-shaped elastic rod deforms in three distinct configurations (deformation modes), presented as Loop 1, Lemniscate, and Loop 2 (fig.(2)). We also determined specific geometrical characteristics of these S-shaped rods leading to these three deformation patterns (table.(2)). This analysis quantitatively determined how specific geometrical parameters of the sagittal spine lead to different deformation patterns of a curved rod, which supported the hypothesis that the variations in the sagittal spinal alignment can a plausible cause of spinal deformity development in adolescent idiopathic scoliosis

The sagittal curvature of the spine in idiopathic scoliosis has been hypothesized to be a mechanical factor leading to the spinal deformity development [27, 28]. However, specific characteristics of the sagittal curve that determines the deformity patterns of the spine are not known. Here, our analysis of S-shaped elastic rods showed that without any assumption about the mechanical properties of the different sections of the spine, geometrical details of the vertebral body or properties of the intervertebral disc, three distinct modes of deformation can be identified for a curved elastic rod under bending and torsion: two loops and one lemniscate. These deformation modes, which are related to the geometrical parameter of the rod (table.(2)), were also observed in the most common scoliotic curve types (Lenke 1 and Lenke 5)[11, 25]. Lenke 1 scoliosis, which manifests as a deformity in the thoracic spine was shown to have axial deformation patterns as seen in Loop 1 and lemniscate. On the other hand Lenke 5 scoliosis had loop shaped deformity patterns as was seen in loop 2 cases. Moreover, the Lenke 1 cases with loop shaped axial projection were shown to have a higher inflection point compared to the Lenke 1 with lemniscate axial projection and Lenke 5 patients[25] (as shown in fig.(5)). This clinical observation matched the simulation results as seen in table.(2). Our reduced order model was able to differentiate between these curve types providing evidence that the elastic rod model as presented here closely reflects the behaviour of the pediatric spine at the onset of scoliosis development.

**Fig. 5:**
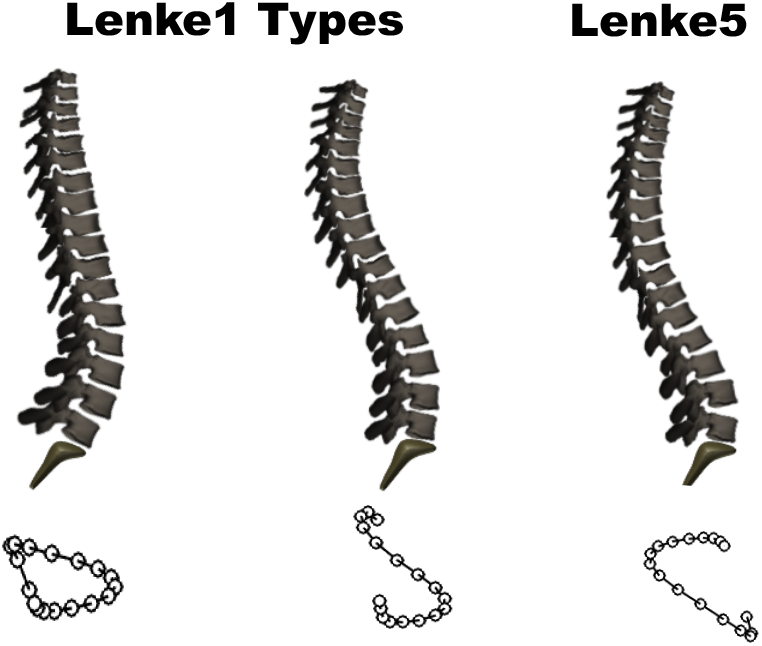
the schematics of the sagittal profiles in Lenke 1 scoliosis with loop (A) and lemniscate (B) axial projection and Lenke 5 scoliosis with loop shaped axial projection.

Fig.(4) and table.(2) show that the average initial configuration is different for the loop and lemniscate cases. *κ_P_* and increase with increase in *θ*_0_ in most cases; exceptions are in lemniscate and Loop2. We see that *κ_P_* ≈ in the three cases. The parameter values are different for the two loop regions. From table.(2), we see that intermediate values of *κ_P_*, and *S* lead to the top view of the deformation being lemniscate shape. Low or high values of these parameters leads to the loop shape. Our results show that we can predict the deformation case based on a limited number of geometrical parameters.

Our analysis further investigate the mechanical loading of the spine in different deformation modes. As shown in fig.(6),(7), although the external loading (gravity and torsion), and mechanical properties of the rods are the same, the moment distribution along the rod varies in the **y-z** and **x-z** planes resulting in different deformation patterns as seen in fig. (8). The geometry affects the final deformations because the moments in the rod depend on *α* and *ϕ* as seen in eq.(7) and (8). *m_y_* causes primarily bending deformations in the **x-z** plane, while *m_z_* causes twisting deformations, bringing the rod out of the **x-y** plane. We see that *m_x_*(*s*) is negative and the rod bends towards **+ve y-**axis in all three cases. *m_x_* changes along the spine in Loop 1, but does not seem to affect the other cases as the deformation in the **y-z** projection can be explained by the effects of *m_x_*(*s*). The reason *m_z_* does not affect the deformation **y-z** projection is due to the difference in magnitude between *m_x_*(*s*) and *m_z_*(*s*).However, when both of these moments are non-zero it is difficult to qualitatively explain the deformation of the rod from a knowledge of their magnitudes. However, as seen in from fig.(6), the shape in the **x-y** projection is predominantly determined by *m_z_*. Note that *m_y_*(*s*) is positive in all three cases of fig.(6), and the axial projection of the three cases follows the same qualitative trend. In particular, the rod deflects towards the **-ve x-**axis for small *s*, and then bend towards the **+ve x-**axis for larger *s* (in the Loop1 case this trend is not as obvious as the other two cases, but the rod begins to bend towards the **+ve x-**axis for larger *s*). Thus, the sign of the *m_y_* moment and the deformation are correlated. The deformed shaped of the sagittal curves are presented in the supplementary materials. It can be seen the relative position of the inflection point between the groups, i.e. loop 1 being the highest and loop 2 the lowest with respect to the horizontal axis as was seen prior to the deformation, holds true in the deformed shape of the spine (Supplementary material fig.(S1))

**Fig. 6:**
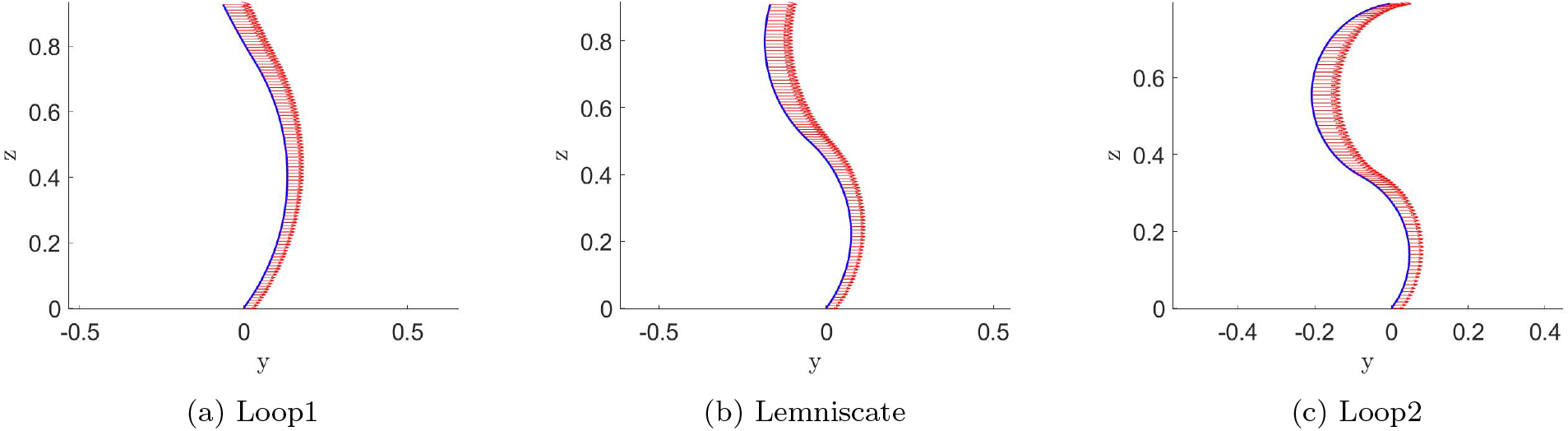
Sagittal view of the initial configuration of the 37° average curve in fig.(4). (*m_y_*(*s*) plotted as red vectors)

**Fig. 7:**
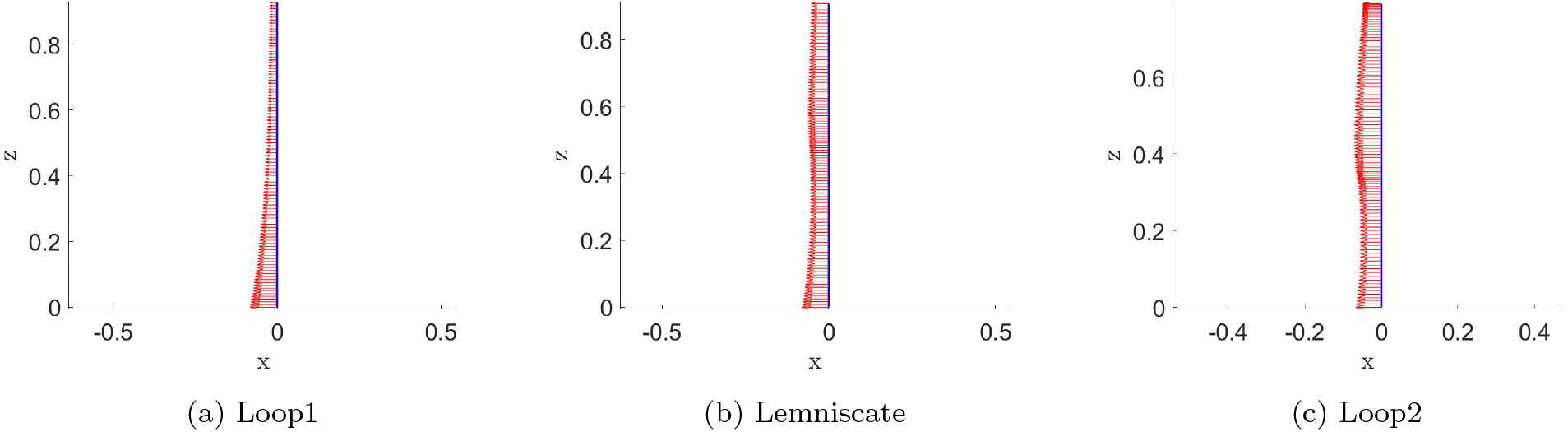
Frontal view of the initial configuration of the 37° average curve in fig.(4). (*m_x_*(*s*) plotted as red vectors)

**Fig. 8:**
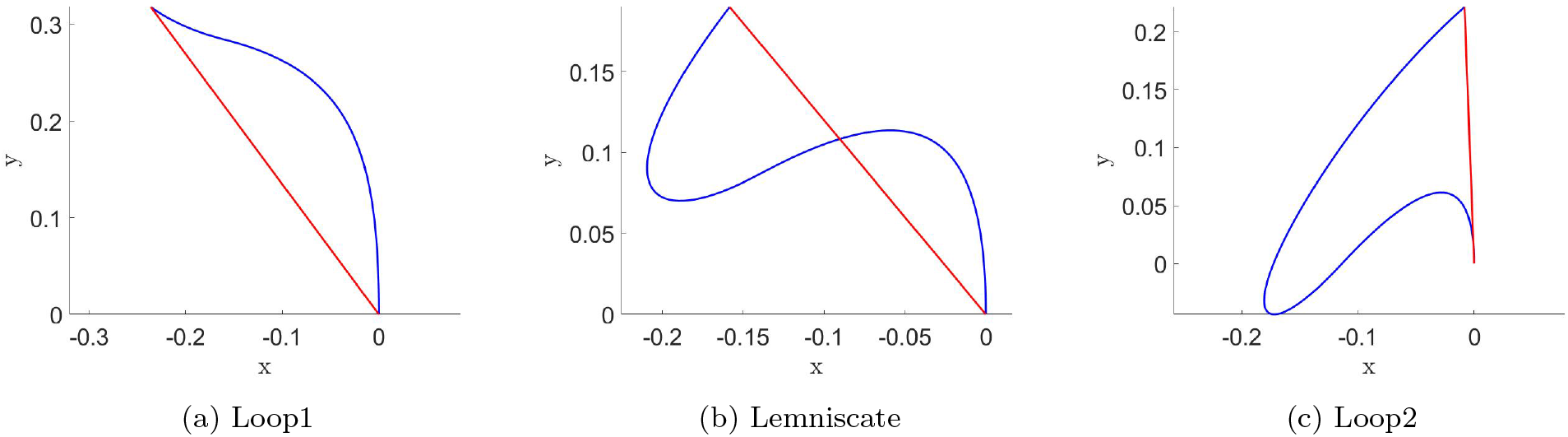
Axial view of the deformed configuration of the 37° average curve in fig.(4). Red line-segment connects end points to show loop/lemniscate classification.

An analysis of the scoliosis deformity patterns, i.e., loop versus lemniscate, are particularly important because they not only help to better understand the pathogenesis of the spinal deformity development in adolescent idiopathic scoliosis but also are shown to impact the clinical management of the disease. In a 3D classification of the spine that aimed to study the bracing effectiveness, a common conservative treatment for small pediatric spinal deformities, it was shown that patients with loop and lemniscate deformation patterns respond differently to bracing[22]. In surgical treatment of idiopathic scoliosis, it was also shown as the lemniscate shape curves consist of two 3D curves as opposed to the loop shaped cases that consist of one 3D curve, the surgical planning of the disease should also distinguish between these curve types [24]. As the presented rod model could determine the loop versus lemniscate shape patterns, future applications of this model in predicting the conservative treatment outcomes or surgical planning of the pediatric spine are warranted.

## 5 Conclusion

By changing the geometry of an S-shaped elastic rod, we determined three different deformation patterns. The deformation patterns were related to the position of the inflection point of the S-shaped rod, and the curvature of the rod above and below the inflection point. These parameters change slightly as the base angle of the rod changes within each deformity pattern group. These curve characteristics relate to the sagittal curvature of the spine in the most common scoliotic curve types. This analysis provided evidence that changes in the sagittal curvature of the spine can be responsible for different deformation patterns in idiopathic scoliosis.

## Acknowledgements

Sunder Neelakantan and Prashant K. Purohit acknowledge partial support for this work through an NSF grant NSF CMMI 1662101.

## Conflict of interest

The authors declare that they have no conflict of interest.

## Notes

### Competing Interest Statement

The authors have declared no competing interest.

